# CRISPR-FOIL: A Programmable CRISPR Tool to Engineer and Illuminate Chromatin Folding in Live Human Cells

**DOI:** 10.64898/2026.08.09.743771

**Authors:** Yu-Chieh Chung, Sydney Willey, Siou-Luan He, Nevin Wise, Li-Chun Tu

**Affiliations:** Department of Biological Chemistry and Pharmacology, The Ohio State University, Columbus, OH, USA 43210; Center for RNA Biology, The Ohio State University, Columbus, OH, USA 43210; The Ohio State University Comprehensive Cancer Center, The Ohio State University, Columbus, OH, USA 43210; Michigan Neuroscience Institute, University of Michigan, Ann Arbor, MI 48109

**Keywords:** CRISPR, chromatin organization, fluorescence microscopy, live-cell imaging

## Abstract

Chromatin organization plays a critical role in regulating gene expression. Chromatin compaction represses gene expression by physically restricting the access of the transcriptional machinery to DNA, while spatial proximity between enhancers and promoters, often mediated by chromatin loops, is essential for gene activation. To investigate the regulatory mechanisms underlying loop formation and chromatin compaction, as well as their effects on gene expression, we developed CRISPR-FOIL (utilizing <u>CRISPR</u> to <u>FO</u>ld and <u>IL</u>luminate chromosomal DNA), a novel programmable platform for engineering chromatin loops and inducing chromatin compaction in live cells. CRISPR-FOIL anchors pairs of genomic loci in proximity by engineered single-guide RNAs (sgRNAs), resulting in an artificial chromatin loop. The fused two CRISPR-Sirius gRNAs enable genomic loci to be visualized through fluorescent RNA coat proteins in various colors. In addition, multiple CRISPR-FOIL complexes can act cooperatively to drive chromatin compaction. These results establish CRISPR-FOIL as a powerful tool for engineering chromatin organization in live cells and highlight its potential as a therapeutic platform for gene regulation and disease control.

## INTRODUCTION

Chromatin organization is crucial for the regulation of gene expression and essential cellular processes, such as cell differentiation, cell cycle progression, and DNA damage repair. In eukaryotes, the establishment of chromatin loops and topologically associating domains (TADs) is mediated by various mechanisms ^1–3^. Chromatin loops mediate transcription by bringing transcription factors into close proximity with promoters, which then elevates the transcription firing rate and efficiency ^4^. For example, chromatin can be looped by structural maintenance of chromosomes (SMC) complexes, such as cohesin, and stabilized by CCCTC-binding factors (CTCF). During interphase, cohesin mediates chromatin looping on a timescale of minutes, contributing to the dynamic structure of chromatin ^5^. Although the disruption of TADs in some somatic cells does not broadly alter gene expression profiles ^6^, TADs play essential roles in normal genome function, including embryonic development and rapid immune responses to environmental stimuli ^7,8^. At a larger scale, groups of loops partition into megabase-scale compartments that segregate transcriptionally active chromatin from inactive regions. However, the establishment and functional impact of these mesoscale chromatin compartments remain largely enigmatic.

Understanding the mechanisms driving dynamic and heterogeneous chromatin organization among cells of the same type remains a major challenge in the field. Although current high-spatial-resolution methods have advanced our understanding, these approaches often rely on static or population-based samples, which obscure temporal dynamics in single cells ^9^. While chromosome conformation capture–based approaches and fluorescence in situ hybridization (FISH) have provided significant insights, they cannot easily distinguish time-dependent fluctuations from cell-to-cell heterogeneity ^10^. Live-cell visualization of chromatin dynamics has been achieved by inserting repetitive bacterial sequences, such as the *lac* operator; however, this strategy is labor-intensive and can introduce structural artifacts into native chromatin ^11^. Alternatively, artificial zinc fingers have been engineered to tether enhancer regions to the b-globin promoter to activate transcription, but this technique requires extensive molecular biology expertise ^12^. Consequently, there is a pressing need for programmable live-cell tools that can simultaneously manipulate (fold) and track (illuminate) long-range chromatin architecture in single cells without introducing structural (functional) artifacts.

The clustered regularly interspaced short palindromic repeats (CRISPR)/Cas system has been repurposed from a bacterial immune defense mechanism into a versatile platform for genome editing, epigenetic regulation, biosensing, and live-cell imaging ^13–15^. The type II CRISPR system from *Streptococcus pyogenes* is widely used due to its precise targeting and high binding affinity ^16^. To streamline applications, the native crRNA and tracrRNA are commonly engineered into a single guide RNA (sgRNA). While catalytically active Cas9 induces double-strand breaks for gene editing, a nuclease-dead variant (dCas9, carrying D10A and H840A mutations) retains strong DNA-binding capability without cleaving DNA, making it ideal for non-editing applications. This CRISPR/dCas9 system has been used to develop live-cell DNA imaging techniques to visualize chromatin organization and track its dynamics ^17^. We previously developed CRISPR-Sirius, a bright and thermostable genome imaging system, to track chromatin dynamics and mesoscale conformations in living cells ^18^.

While earlier efforts induced chromosomal looping within sub-kilobase ranges to activate transcription, those approaches were restricted to regulatory elements across short genomic distances ^19^. A programmable tool to manipulate and visualize long-range chromatin loops and mesoscale architecture in live cells is still lacking. To address this gap, we expanded CRISPR-Sirius into CRISPR-FOIL (CRISPR to both FOld and ILluminate chromosomal DNA), a dual-functional platform capable of simultaneous chromatin looping and real-time imaging across kilobase to megabase scales. Beyond stabilizing individual chromatin loops, CRISPR-FOIL can target multiple regions along a single chromosome to induce mesoscale chromosome contraction. Ultimately, CRISPR-FOIL enables direct manipulation of chromatin architecture for functional studies in living cells, while opening new avenues for therapeutic and diagnostic interventions targeting pathogenic extrachromosomal DNA (ecDNA) driving cancer progression.

## RESULTS

### Engineering and imaging stable *de novo* chromatin loops using CRISPR-FOIL guide RNAs

CRISPR-Sirius single-guide RNAs (Sirius-gRNAs)^18^ carry an octet of stem-loops from bacteriophages (e.g., MS2 or PP7) that bind fluorescently labeled RNA coat proteins, enabling the fluorescent labeling of specific genomic regions (Figure 1A, top). CRISPR-Sirius has been successfully used to investigate the dynamics of specific genomic loci and mesoscale chromatin conformations with high efficiency and specificity ^10,20^. These results laid the foundation for developing CRISPR-FOIL, a programmable platform for engineering chromatin looping and studying human genome function.

**Figure 1.**
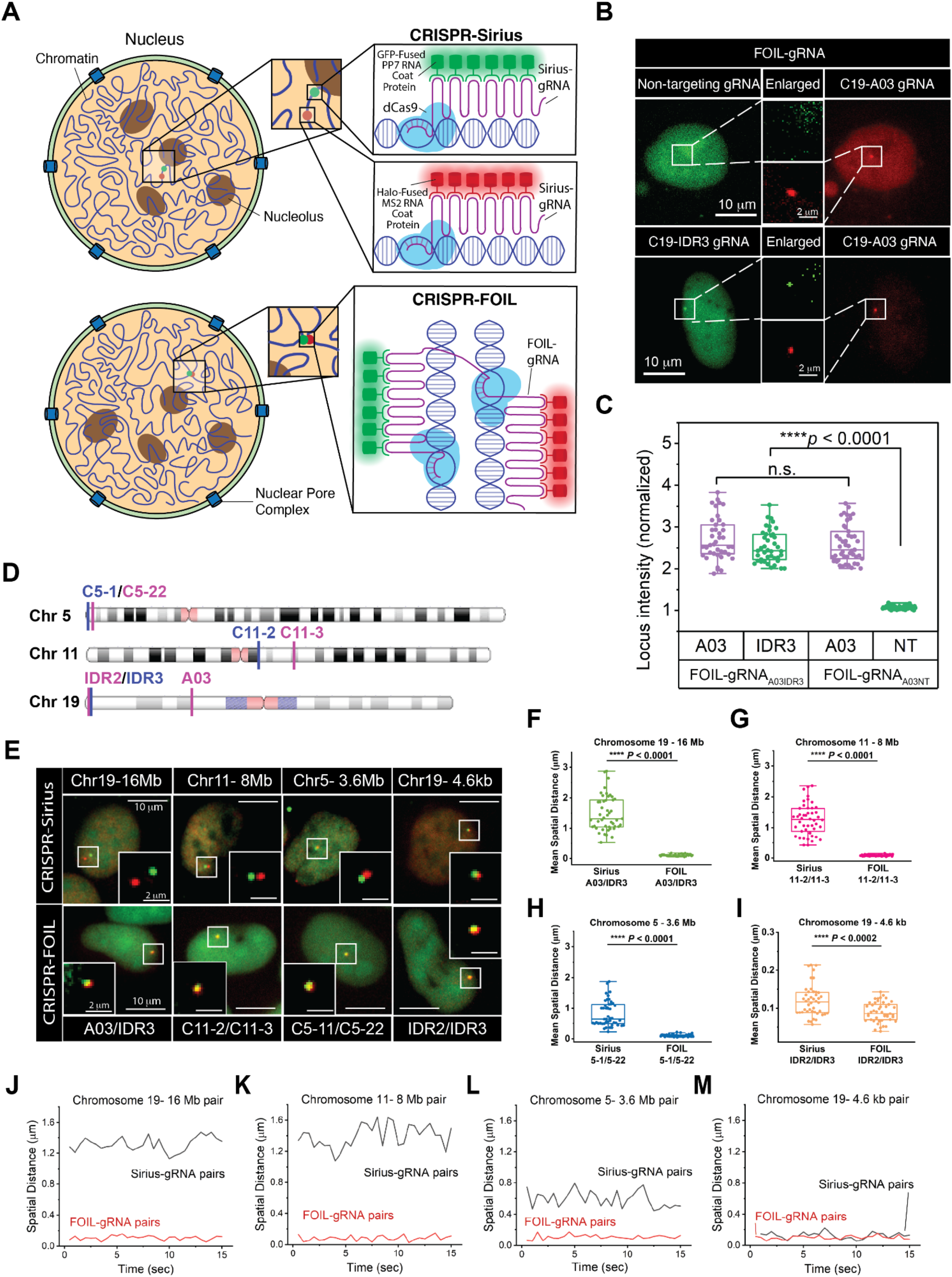
CRISPR-FOIL drives chromatin looping. (A) Schematic representation of CRISPR-Sirius (top) and CRISPR-FOIL (bottom). CRISPR-FOIL adopts the design of CRISPR-Sirius, enabling the targeting and visualization of specific loci, as well as bringing targeted loci into proximity by pairing different sgRNA stem loops (FOIL-sgRNA). (B) Representative images of CRISPR-FOIL–targeted loci. Specificity of CRISPR-FOIL targeting by pairing a non-target sgRNA with the C19-A03 sgRNA (top), and by pairing two different sgRNAs targeting the C19-IDR3 and C19-A03 loci (bottom). (C) Box plot of local fluorescence intensity of targeted loci shown in (B). N_C19-A03-Non-target-FOIL_=47, N_C19-A03-C19-IDR3- FOIL_=41. (D) Genomic locations of CRISPR-FOIL–targeted loci on Chr 5, Chr 11and Chr 19. (E) Representative images of loci targeted by CRISPR-FOIL and CRISPR-Sirius.at the locations shown in (D). (F) Box plot of distances between the pair of loci C19-A03 and C19-IDR3 targeted by CRISPR-FOIL and CRISPR-Sirius. N_C19-A03-C19-IDR3-FOIL_=41, N_C19-A03-C19-IDR3-Sirius_=47. (G) Box plot of distances between the pair of loci C11-2 and C11-3 targeted by CRISPR-FOIL and CRISPR-Sirius. N_C11-2C11-3--FOIL_=45, N_C11-2C11-3-Sirius_=46. (H) Box plot of distances between the pair of loci C5-1 and C5-22 targeted by CRISPR-FOIL and CRISPR-Sirius. N_C5-1C5-22--FOIL_=41, N_C5-1C5-22--Sirius_=47. (I) Box plot of distances between the pair of loci C19-IDR2 and C19-IDR3 targeted by CRISPR-FOIL and CRISPR-Sirius. N_C19- IDR2-C19-IDR3-FOIL_=45, N_C19-IDR2-C19-IDR3-Sirius_=44. (J)–(M) Representation of the temporal changes in spatial distances between pairs of loci targeted by CRISPR-FOIL and CRISPR-Sirius. Imaging was performed over 30 frames at 500 ms intervals. All experiments were repeated at least three times.

To engineer and visualize *de novo* chromatin looping using a single Cas system, we redesigned the sgRNAs as chimeric constructs (FOIL-gRNAs) by joining two distinct Sirius-gRNAs that target separate genomic regions on the same chromosome (Figures 1A bottom and S1A). RNAfold web server simulations predicted that these FOIL-gRNAs fold into stable secondary structures (Figure S1B). To express full-length FOIL-gRNAs, we used a Pol II promoter (CMV) instead of the Pol III promoter (U6) used for standard Sirius-gRNA expression, as the latter resulted in poor expression of fused Sirius-gRNAs (Figure S2).

We first tested the targeting specificity of FOIL-gRNA using one non-targeting and one targeting sgRNA pair, alongside a pair containing two targeting sgRNAs as a control. Only targeted loci produced fluorescent signals, with no signal observed in the non-target channel (Figure 1B). Quantitative analysis of fluorescence intensity confirmed the targeting specificity of CRISPR-FOIL (Figure 1C, Materials and Methods). Specifically, our data demonstrate that FOIL-gRNA with both targeting sequences yields significantly higher signals across both channels, whereas FOIL-gRNA with only one targeting sequence produces signals predominantly in its corresponding targeting channel (Figures 1B and 1C).

We next applied CRISPR-FOIL to engineer stable chromatin loops across different chromosomes and varying genomic distances between locus pairs. Target sites were selected on chromosomes 5, 11, and 19, each possessing distinct gene densities and preferential nuclear localizations reported by previous studies^19^. Chromosome 19 (Chr19) is gene-rich and centrally localized; chromosome 11 (Chr11) is gene-rich and peripherally localized; whereas chromosome 5 (Chr5) is relatively gene-poor and primarily localized at the nuclear periphery.

To test the spatial extent of CRISPR-FOIL looping, target genomic loci were paired across varying linear distances: (1) C5-1 and C5-22, 3.6 Mb apart on the Chr5 p-arm; (2) C11-2 and C11-3, 8 Mb apart on the Chr11 q-arm; (3) A03 and IDR3, 16 Mb apart on the Chr19 p-arm; and (4) IDR2 and IDR3, 4.6 kb apart on the Chr19 p-arm (Figure 1D and Table S1). These loci were successfully targeted by both CRISPR-Sirius and CRISPR-FOIL (Figure 1E). The successful targeting of individual gRNAs was also validated by pulling down dCas9-bound chromatin via ChIP-PCR (Figure S3). Imaging and quantification of the spatial distances between loci labeled by FOIL-gRNAs revealed significantly closer and partially overlapping signals compared to those labeled by Sirius-gRNAs, indicating that CRISPR-FOIL successfully generated chromatin loops in live cells (Figures 1E and 1F–1I). Specifically, locus pairs separated by 3 to 16 Mb along linear DNA typically exhibited 3D spatial distances of 500 nm to 2 µm in the nucleus; with CRISPR-FOIL, this distance was reduced to approximately 100 nm. Even for the short-range 4.6 kb IDR2/IDR3 pair, CRISPR-FOIL significantly reduced spatial distance compared to CRISPR-Sirius (*p* < 0.0002).

Finally, we tracked real-time distance fluctuations between locus pairs under CRISPR-FOIL (looped) and CRISPR-Sirius (unlooped) conditions. Loci linked by CRISPR-FOIL exhibited constrained movement compared to the larger fluctuations seen with CRISPR-Sirius, highlighting the stability of CRISPR-FOIL– mediated looping (Figures 1J–1M and S4). Together, these results demonstrate that CRISPR-FOIL is a programmable platform for engineering stable, *de novo* chromatin loops across a wide range of genomic distances.

### CRISPR-FOIL-mediated cross-linking drives mesoscale chromosome contraction and represses gene transcription

Genes within constitutive heterochromatin, marked by H3K9me3 are largely repressed through cross-linking by heterochromatin protein 1 (HP1) dimers ^21^. To investigate whether mesoscale chromosomal silencing could be achieved by cross-linking multiple genomic regions within a single chromosome using CRISPR-FOIL, we designed FOIL-gRNA to target approximately 836 copies of repetitive sequences on the q arm of chromosome 19 (C19q) (Figure 2A and Table S1), spanning a ∼17 Mb region across half of the arm and excluding the pericentromeric region. The C19q targeting region was previously verified in U2OS cells using FISH oligopaint ^22^. Given that the entire q arm encodes roughly 994 genes (of which approximately 594 are active)^10^, it is reasonable to estimate that more than a hundred active genes reside within the targeted C19q region, making it a good model system to evaluate the link between cross-linking-induced chromatin compaction and transcriptional output.

**Figure 2.**
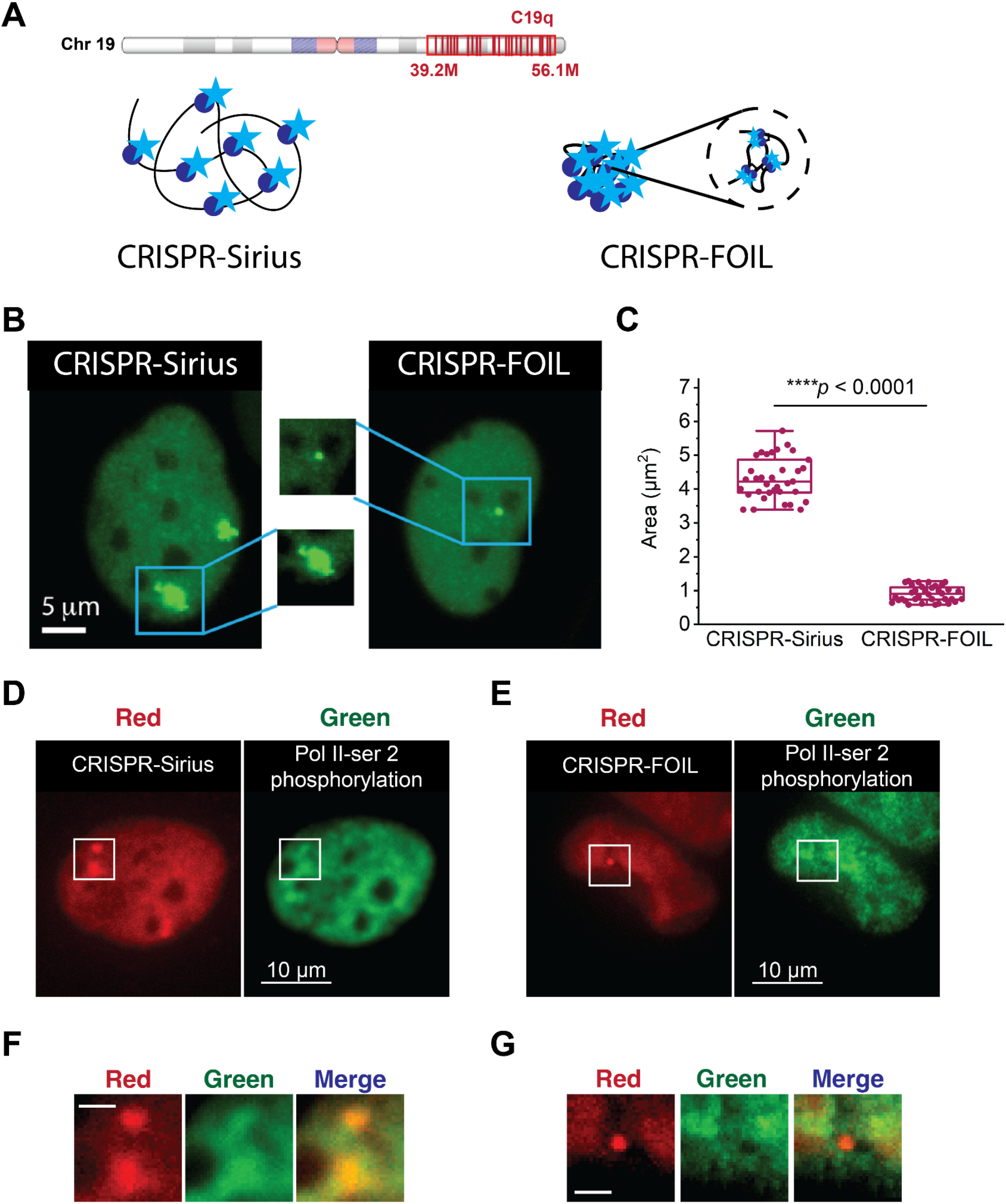
Multiple FOIL-gRNAs cooperatively induce mesoscale chromatin contraction and decrease transcription. (A) Schematic of mesoscale chromatin (C19q) contracted by CRISPR-FOIL. Top: Genomic targeting of C19q, spanning ∼17 Mb on the chr19 q-arm. Bottom left: Schematic of C19q labeled with CRISPR-Sirius sgRNAs without inducing artificial cross-linking. Bottom right: Schematic of C19q-FOIL-sgRNAs simultaneously crosslinking multiple distinct targeting sites (∼836) within chr19 q- arm to induce C19q contraction. (B) Representative images of mesoscale chromatin (C19q) labeled by CRISPR-Sirius sgRNAs (left) or multiple FOIL-sgRNAs (right). (C) Boxplot of C19q area in cells targeted by C19q-Sirius-sgRNAs (N_C19q-Sirius_ =35) vs. C19q-FOIL-sgRNAs (N_C19q-FOIL_=36). (D) Representative image of C19q labeled by CRISPR-Sirius sgRNAs (left) and corresponding immunofluorescence staining against RNA polymerase II serine 2 phosphorylation (Pol II-Ser2P, right). (E) Representative image of C19q labeled by C19q-FOIL-sgRNAs (left) and corresponding Pol II-Ser2P immunofluorescence staining (right). (F) Enlarged image of C19q from (D, left), corresponding Pol II- Ser2P signal from (E, right), and merged overlay (right). (G) Enlarged image of C19q from (E, left), corresponding Pol II-Ser2P signal from (E, right), and merged overlay (right). Scale bars in (F) and (G) represent 2 µm. Pseudo-colors were applied to images for better visualization. All experiments were repeated at least three times.

Strikingly, CRISPR-FOIL induced a ∼5-fold reduction in C19q area compared to CRISPR-Sirius controls (Figure 2B and 2C). To test whether this physical contraction coincides with transcriptional repression, we performed immunofluorescence staining against RNA polymerase II serine 2 phosphorylation (Pol II-pSer2), a post-translational modification associated with active transcriptional elongation. By analyzing the overlap between the C19q loci in a double red U2OS line (U2OS^dCas9-HSA/PCP-tdTOMATO/MCP-HaloTag^, see Material and Methods for details) and Pol II-pSer2 (labeled with green, Alexa 488) fluorescence signals, we observed that C19q labeled with Sirius-gRNA overlapped with regions exhibiting higher Pol II-pSer5 intensity (Figures 2D and 2F), indicating that transcription remained active. This finding is consistent with our previous qPCR results in CRISPR-Sirius-labeled cells ^20^. Conversely, the contracted C19q region in CRISPR-FOIL cells localized to areas with lower Pol II-pSer2 signals (Figures 2E and 2G), implying a reduction in transcriptional activity. Together, these results demonstrate that synthetic cross-linking via CRISPR-FOIL drives mesoscale chromatin condensation and localized gene suppression.

## DISCUSSION

Although various tools have been developed to manipulate chromatin organization, most act broadly by altering histone modifications across the genome. For example, small epigenetic inhibitors targeting methyltransferases and deacetylases are widely used to induce global chromatin reorganization and reverse epigenetic dysregulation in cancer^23^. However, live-cell tools capable of locally manipulating chromatin loops and compartments are only beginning to emerge. Here, we demonstrated that CRISPR- FOIL can selectively tether specific genomic loci to induce targeted artificial chromatin looping. By employing FOIL-sgRNAs designed against low-repetitive sequences, we achieved reliable targeting while maintaining robust fluorescence signal intensity for tracking.

While engineering loops at non-repetitive genomic regions remains challenging with single sgRNA constructs, CRISPR-FOIL may be used to overcome this limitation when combined with a multi-sgRNA strategy targeting adjacent sites within narrow genomic windows. This multi-sgRNA strategy approach significantly enhances the signal-to-noise (S/N) ratio and likely generates sufficient localized mechanical force to drive stable loop formation. Moreover, when targeted to highly repetitive regions, CRISPR-FOIL promotes cooperative, massive cross-linking that compacts mesoscale chromatin and represses transcriptional output. Together, these features establish CRISPR-FOIL as a versatile, programmable platform to manipulate chromatin architecture across diverse length scales and investigate its functional consequences in living cells.

Looking ahead, CRISPR-FOIL offers strong translational potential for studying and treating pathologies linked to higher-order genome misregulation. Beyond restoring disrupted enhancer–promoter interactions, this technology could be leveraged to compact and silence pathological genomic features, such as extrachromosomal DNA (ecDNA) or supernumerary chromosomes. Furthermore, incorporating single nucleotide polymorphisms (SNPs) into sgRNA design will enable allele-specific chromatin manipulation, enhancing target precision while avoiding off-target contraction of essential homologous alleles to minimize cellular toxicity.

## MATERIALS AND METHODS

### Plasmid construction

We genetically fused Sirius-sgRNA-8XPP7 (pPUR-sgRNA-Sirius-8XPP7, Addgenet#121940) and Sirius-sgRNA-8XMS2 (pPUR-sgRNA-Sirius-8XMS2, Addgene#121939) with a synthesized linker (2xBSL)^24^ to make pPUR-hU6-sgRNA-8XMS2-2xBSL-sgRNA-8XPP7, in which the fused Sirius- sgRNA is driven by a human U6 promoter (hU6). Due to poor gRNA expression, we then switched the hU6 promoter (pol III) to a CMV (pol II) promoter^25^ from pHAGE-TO-dCas9-P2A-HSA (Addgene#121936) by PCR. Two ribozymes were added in the plasmid - the hammerhead (HH) ribozyme to the 5’ end and hepatitis delta virus (HDV) ribozyme at the 3’ end of the gRNA to remove extra nucleotides generated during pol II transcription at both ends (i.e., cap and polyA tail). The resulting construct is named pPUR-CMV-FOIL1-gRNA-8XMS2-2XBSL-FOIL2-gRNA-8XPP7. The sgRNA targeting sequences, CGATGAGAGGCCnGG for C5-1, TGTGTCTTGCACnGG for C5-22, CACACATCAACAnGG for C11-2, CTAGGACTCAGAnGG for C11-3, CCnCCCAGCCTCTCC for C19-IDR2, AGCAGATGTAGGnGG for C19-IDR3, AATGAGTGAGCGnGG for C19-A03, and TCCCTCAGACCCnGG for C19q (Table S1), were purchased from Integrated DNA Technologies, and cloned into the sgRNA vector using BbsI sites (Figure S2). The cloned sgRNA constructs were confirmed by Sanger sequencing. Constructs for dCas9, GFP-fused PP7 coat protein, and HaloTag-fused MCP coat protein expression were obtained from Addgene (#121936, #121938, and #121937, respectively, Table S2). Designs of these plasmids can be found in previous work^18,26^. pHAGE-EFS-PCP-tdTOMATOnls was made by replacing GFP to a codon optimized tdTOMATO in pHAGE-EFS-PCP-GFPnls by using BamHI and NotI sites (see supplementary Information for its sequence).

### Cell culture

The wild-type human bone osteosarcoma epithelial (U2OS) cell line was a kind gift from Dr. Thoru Pederson’s laboratory. The dual color U2OS line U2OS^dCas9-HSA/PCP-GFP/MCP-HaloTag^ was established in previous work^18^. The human embryonic kidney epithelial (HEK293T) cells were obtained from ATCC. U2OS cells and stable derivatives were maintained in DMEM with high glucose, supplemented with 10% FBS and 1% penicillin/streptomycin. HEK293T cells were maintained in Iscove’s modified Dulbecco’s medium supplemented with 1% GlutaMAX, 10% FBS and 1% penicillin-streptomycin. All cell lines were routinely tested and confirmed to be free of mycoplasma contamination using a Mycoalert PLUS kit (Lonza).

### Lentiviral transduction

To establish stable cell lines with fluorescently labeled C19q, lentiviral particles carrying the sgRNA plasmids were generated in HEK293T cells as previously described^27^. Approximately 5×10^6^ cells were seeded in 6 well plates 24 hours before transfection. Cells were cotransfected with 0.5µg of pCMV-dR8.2 dvpr (Addgene), 0.3µg of pCMV-VSV-G (Addgene), and 1.5 µg of the sgRNA plasmid were using TransIT transfection reagent (Mirus) according to the manufacturer’s protocol. 48 hours post-transfection, the virus was collected and filtered through a 0.45µm filter, and either used immediately or snap frozen and stored at -80°C. The U2OS cells were transduced by spinfection in 6-well plates. Approximately 2×10^5^ cells were incubated with 1 ml of lentiviral supernatant and centrifuged at 1,200 x g for 30 minutes.

### Flow cytometry

We established a double red U2OS line, U2OS^dCas9-HSA/PCP-tdTOMATO/MCP-HaloTag^, for C19q experiments, in which MCP-HaloTag and the red PCP (PCP-tdTomato) double positive cells were sorted (Figure S5). The U2OS cells were first transduced to express dCas9 and MCP-HaloTag and selected using Alexa647- conjugated anti-mouse CD24 antibody (BioLegend) and HaloTag-JF549 (Promega). A final concentration of 2 nM HaloTag-JF549 was added 12-16 hours before cell sorting, and HSA was stained following the manufacturer’s staining protocol. Next, PCP-tdTomato was transduced into the sorted U2OS cells U2OS^dCas9-HSA/MCP-HaloTag^ and PCP-tdTomato positive cells were sorted while leaving the MCP-HaloTag unstained. Sorting was performed on a BD fluorescence-activated sorter (FACSAria Fusion cell sorter) equipped with 405-, 488-, 561-, and 637-nm excitation lasers. MCP-HaloTag-JF549 and PCP-tdTomato were detected with a 575/26 nm emission filter, and dCas9 positive cells were detected with a 670/30 nm emission filter.

### Chromatin Immunoprecipitation-PCR

The ChIP assays were performed using U2OS C19-A03/IDR3, U2OS C5-1/C5-22, U2OS C11-2/C11-3, and U2OS C19-IDR2/IDR3 cells. 4×10^6^ cells were collected per cell line, and protein and DNA were cross-linked in 1% formaldehyde at room temperature for 10 minutes. The cross-linking reaction was then quenched by adding 2.5 M glycine to a final concentration of 0.125 M, followed by incubation on ice for 5 minutes. After washing the fixed cell pellet, the chromatin was sheared to an average size of 200 bp using a Covaris sonicator. A small aliquot of the supernatant was used as the input control, and the remaining sonicated chromatin was divided into two aliquots for incubation with either an anti-Cas9 antibody (Takara, 632607) or an anti-IgG antibody (SouthernBiotech, 0111-01) as a negative control. The antibody-chromatin complexes were precipitated with Dynabeads Protein A (Invitrogen). DNA fragments were released from the immunoprecipitated complexes by reversing the cross-linking at 65 °C overnight. The precipitated DNA was purified using the MinElute PCR Purification Kit (QIAGEN) and used as a template for PCR.

### Immunofluorescence Staining

In a 35 mm glass-bottom imaging dish (MetTek), approximately 1×10^5^ cells were fixed in 1% paraformaldehyde at room temperature for 15 minutes in cytoskeletal buffer (10 mM PIPES, 300 mM sucrose, 100 mM NaCl, 2 mM MgCl_2,_ 1mM EGTA, pH 6.8). The cells were then washed with TBS-1 buffer (10 mM Tris-HCl, 150 mM NaCl, 3 mM KCl, 1.5 mM MgCl_2_, 0.2% glycine, 0.05% Tween 20, 0.1% BSA, pH 7.7) and permeabilized in cytoskeletal buffer containing 0.5% Triton X-100 at room temperature for 30 minutes. Following permeabilization, the cells were washed three times with TBS-1 buffer and blocked in TBS-1 at room temperature for 30 minutes. Cells were then incubated with a primary antibody against human RNA Polymerase II serine 2 phosphorylation (Active Motif, 61083) at 4 °C for 12 hours. After three subsequent washes with TBS-1, cells were incubated with a secondary goat anti-rat IgG (H+L) Alexa Fluor 488 antibody (ThermoFisher, A11006) for 30 minutes at room temperature. Finally, the cells were washed three times with TBS-1 and mounted using ProLong Gold antifade reagent prior to imaging.

### Fluorescence microscopy and image processing

A custom-built Olympus IX83 microscope equipped with three EMCCD cameras (Andor iXon 897), a LED, 60X apochromatic oil objective lens (NA 1.5), and mounted with a 1.6X magnification adaptor, resulting in a total magnification of 96X. The microscope incubation chamber was maintained at 37°C supplied with 5% CO2 v/v and humidity for live cell imaging. Image data were acquired using CellSens software (version 4.1.1). For C19q imaging, z-slices per nucleus were acquired with an exposure time of 100 ms: 31 slices with a 0.22 µm step size for live U2OS cells, and 15 slices with a 0.15 µm step size for fixed U2OS cells. The images of a z-stack series were projected to a 2D image by using maximum intensity projection. The brightness and contrast of images were then adjusted to the best visibility.

Images were registered and analyzed using Fiji ^28^ and Mathematica (Wolfram) software. Images obtained by using green and red channels were registered by 0.1 µm coverglass-absorbed TetraSpeck fluorescent microspheres (Invitrogen) as a standard sample. Intensity ratios in Figure 1C were calculated using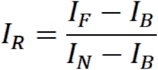 where *I*_*R*_ is the intensity ratio between the labeled loci (*I*_*F*_) and the nucleoplasm (*I*_*N*_). The background fluorescence intensity (*I*_*B*_) from a dark region in the same image was subtracted. The specific genomic locus signals were identified and tracked via the TrackMate plug-in^29^. The spatial distance of locus pairs was calculated by Mathematica, and graphs were generated by OriginPro (OriginLab).

### Analysis of the distance of pairs of loci and temporary fluctuations

The mean spatial distance <R> of a locus pair A and B was calculated by using the time average <R(t)>_t_ of the spatial distance |P_A_(t)-P_B_(t)| over 30 time frames where P(t) is the position vector of a locus at time t. The temporal fluctuation ^δ^R(t) of the spatial distance of a locus pair was calculated by the absolute deviation of spatial distance at time t from the time average of the spatial distance, ^δ^R(t)=|R(t)− <R(t)>_t_|. The mean distance fluctuation ^δ^R was calculated by using the time average < ^δ^R(t)>_t_ (Figure S4).

## AUTHOR CONTRIBUTIONS

Conceptualization, LCT; Methodology, YCC and LCT; Data analysis, YCC performed biophysical analysis; Investigation, YCC, SLH, and LCT performed experiments. Resources, LCT, SW, and NW made the stable cell lines; LCT wrote original draft and made figures; Manuscript reviewing and editing, all authors contributed; Funding acquisition and project supervision, LCT.

## ACKNOWLEDGEMENT

We thank Jenna Thuma and Dilshodbek Nishonov for early work on this project. The authors also acknowledge Dr. Michael Poirier and the Tu lab members for their insightful discussions and comments. This work is supported by the NIH grants R35 GR132998, and OSU start-up fund to L-C.T.

## REFERENCE

1. Kim, Y., Shi, Z., Zhang, H., Finkelstein, I. J. & Yu, H. Human cohesin compacts DNA by loop extrusion. Science 366, 1345–1349 (2019).

2. Davidson, I. F. et al. DNA loop extrusion by human cohesin. Science 366, 1338–1345 (2019).

3. Zuin, J. et al. Nonlinear control of transcription through enhancer–promoter interactions. Nature604, 571–577 (2022).

4. Haberle, V. & Stark, A. Eukaryotic core promoters and the functional basis of transcription initiation. Nat. Rev. Mol. Cell Biol. 19, 621–637 (2018).

5. Rao, S. S. P. et al. Cohesin loss eliminates all loop domains. Cell 171, 305–320.e24 (2017).

6. Schwarzer, W. et al. Two independent modes of chromatin organization revealed by cohesin removal. Nature 551, 51–56 (2017).

7. Stik, G. et al. CTCF is dispensable for immune cell transdifferentiation but facilitates an acute inflammatory response. Nat. Genet. 52, 655–661 (2020).

8. Rajderkar, S. et al. Topologically associating domain boundaries are required for normal genome function. Commun. Biol. 6, 435 (2023).

9. Dekker, J. et al. Spatial and temporal organization of the genome: Current state and future aims of the 4D nucleome project. Mol. Cell 83, 2624–2640 (2023).

10. Chung, Y.-C., Bisht, M., Thuma, J. & Tu, L.-C. Single-chromosome dynamics reveals locus-dependent dynamics and chromosome territory orientation. J. Cell Sci. 136, (2023).

11. Amiad-Pavlov, D. et al. Live imaging of chromatin distribution reveals novel principles of nuclear architecture and chromatin compartmentalization. Science Advances 7, 6251–6253 (2021).

12. Deng, W. et al. Controlling long-range genomic interactions at a native locus by targeted tethering of a looping factor. Cell 149, 1233–1244 (2012).

13. Park, J. S. et al. Digitized kinetic analysis enhances genotyping capacity of CRISPR-based biosensing. ACS Nano 18, 18058–18070 (2024).

14. Pickar-Oliver, A. & Gersbach, C. A. The next generation of CRISPR-Cas technologies and applications. Nat. Rev. Mol. Cell Biol. 20, 490–507 (2019).

15. Thuma, J.Chung, Y.-C. & Tu, L.-C. Advances and challenges in CRISPR-based real-time imaging of dynamic genome organization. Front Mol Biosci 10, 1173545 (2023).

16. Jiang, F., Zhou, K., Ma, L., Gressel, S. & Doudna, J. A. A Cas9–guide RNA complex preorganized for target DNA recognition. Science 348, 1477–1481 (2015).

17. Kim, J. H. et al. LADL: light-activated dynamic looping for endogenous gene expression control.Nature Methods 2019 16:716, 633–639 (2019).

18. Ma, H. et al. CRISPR-Sirius : RNA scaffolds for signal amplification in genome imaging. Nat.Methods 15, (2018).

19. Mehta, I. S., Kulashreshtha, M., Chakraborty, S., Kolthur-Seetharam, U. & Rao, B. J. Chromosome territories reposition during DNA damage-repair response. Genome Biol. 14, R135 (2013).

20. Bisht, M. et al. Differential regulation of mesoscale chromosome conformations in osteoblasts and osteosarcoma. Genome Biol. 26, 307 (2025).

21. Hiragami-Hamada, K. et al. Dynamic and flexible H3K9me3 bridging via HP1β dimerization establishes a plastic state of condensed chromatin. Nat. Commun. 7, 11310 (2016).

22. Feng, Y. et al. Simultaneous epigenetic perturbation and genome imaging reveal distinct roles of H3K9me3 in chromatin architecture and transcription. Genome Biol. 21, 296 (2020).

23. Yu, X. et al. Cancer epigenetics: from laboratory studies and clinical trials to precision medicine.Cell Death Discov. 10, 28 (2024).

24. Hatch, S. C. et al. Gag-dependent enrichment of HIV-1 RNA near the uropod membrane of polarized T cells. J. Virol. 87, 11912–11915 (2013).

25. Xu, L., Zhao, L., Gao, Y., Xu, J. & Han, R. Empower multiplex cell and tissue-specific CRISPR-mediated gene manipulation with self-cleaving ribozymes and tRNA. Nucleic Acids Res. 45, 1–9 (2017).

26. Ma, H. et al. Multiplexed labeling of genomic loci with dCas9 and engineered sgRNAs using CRISPRainbow. Nat. Biotechnol. 1–4 (2016).

27. Ma, H. et al. CRISPR-Cas9 nuclear dynamics and target recognition in living cells. J. Cell Biol.jcb.201604115 (2016).

28. Schindelin, J. et al. Fiji: an open-source platform for biological-image analysis. Nat. Methods 9, 676–682 (2012).

29. Tinevez, J.-Y. et al. TrackMate: An open and extensible platform for single-particle tracking. Methods 115, 80–90 (2017).

