## Supplementary Information for "CRISPR-FOIL: A Programmable CRISPR Tool to Engineer and Illuminate Chromatin Folding in Live Human Cells"

**This PDF file includes:**

> **Figures S1 to S5**
> **Supplementary Table 1 to 2**
> **pHAGE-EFS-PCP-NLSb-tdTOMATOnls sequence**

### SUPPLEMENTARY INFORMATION

**A**

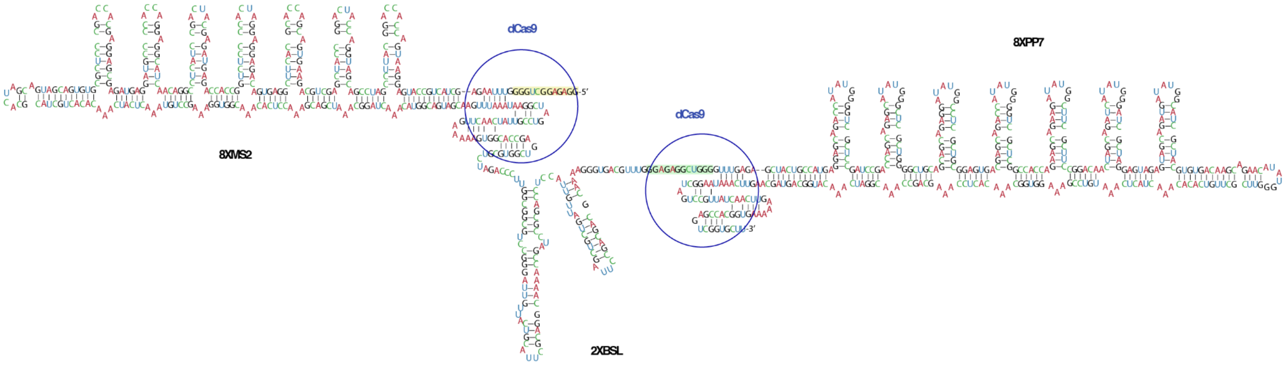


**B**

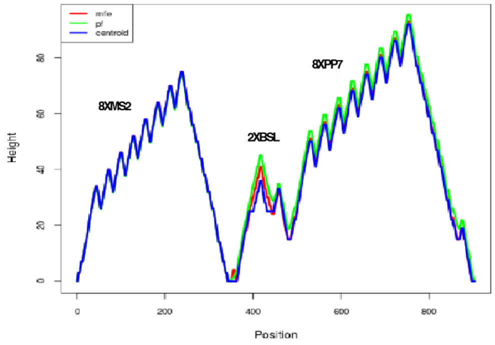


**Supplementary Figure 1**

**Design of CRISPR-FOIL gRNA scaffold.** (A) The diagram depicts the design of CRISPR-FOIL gRNA scaffold. Two CRISPR Sirius-sgRNA (CRISPR-Sirius-sgRNA-8XMS2 and CRISPR-Sirius-sgRNA-8XPP7) were fused, with a linker containing 2xBSL stem loops to enhance the stability of FOIL-gRNA. Two dCas9 binding sites are shown as blue circles. The C19-IDR3 targeting sequence is highlighted in yellow and the C19-IDR2 targeting sequence is highlighted in green. (B) The mountain plot of (A), generated by using RNAfold (version 2.6.3), illustrates the thermodynamic ensemble of the RNA structure, highlighting changes in free energy across the RNA sequence. Structural stability is highest at nucleotide 331 and 896, where maximal base pairing occurs. In contrast, the lowest stability (highest free energy) peaks at the nucleotide 240 and 755, where the eighth MS2 and PP7 stem loop begins to form base pairs. Overall, CRISPR-FOIL gRNA structure is thermostable in its full length.

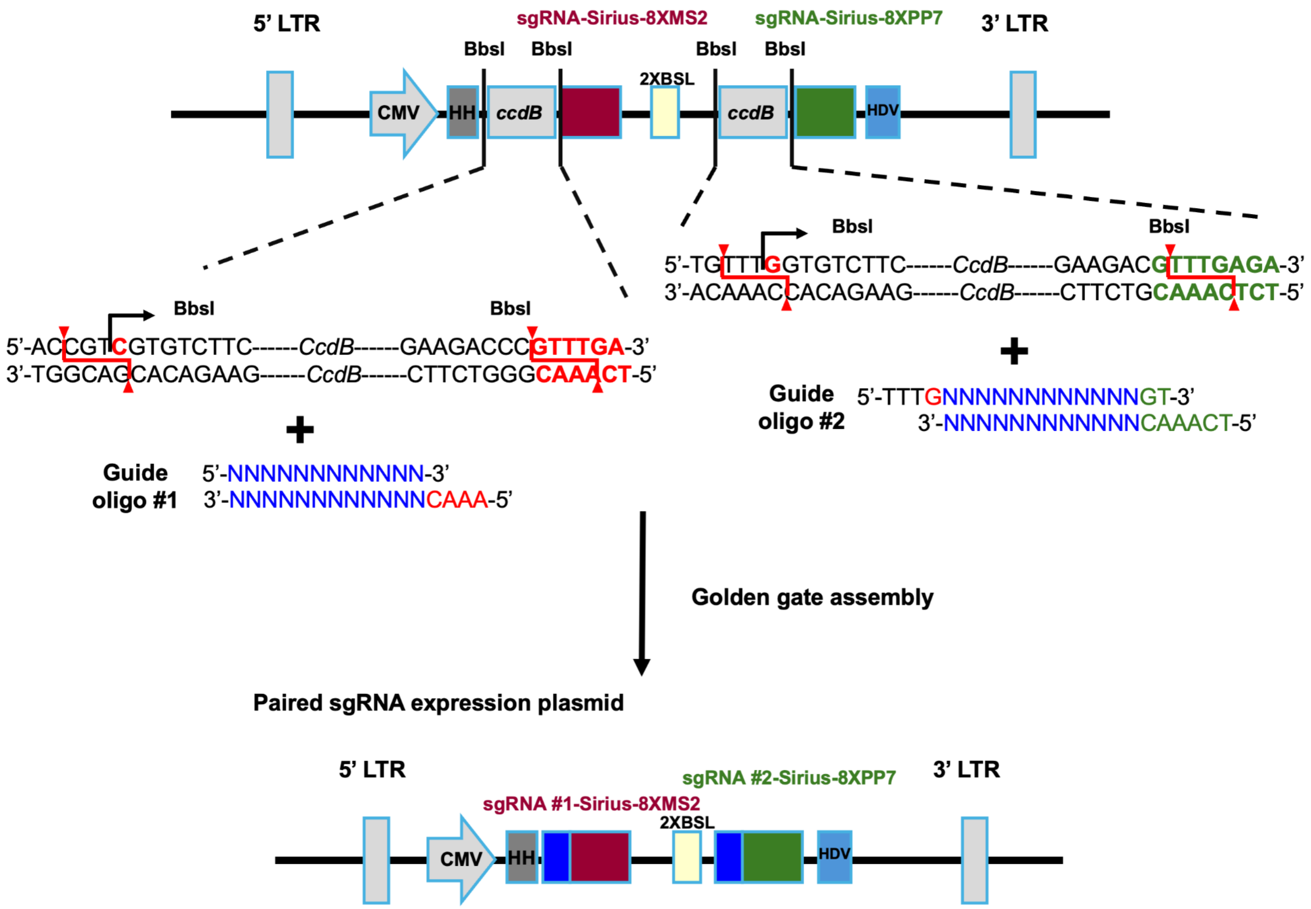


**Supplementary Figure 2**

**Single-step cloning for inserting two FOIL gRNA into the CRISPR FOIL-gRNA construct.** The restriction enzyme BbsI is used to remove two ccdB elements, allowing the insertion of sgRNA sequences targeting specific genomic regions. The sgRNA sequences, synthesized commercially with distinct overhanging sticky ends, were annealed into double-stranded DNA using a PCR machine. Competent cells containing undigested vectors will not grow due to the toxicity of the ccdB gene.

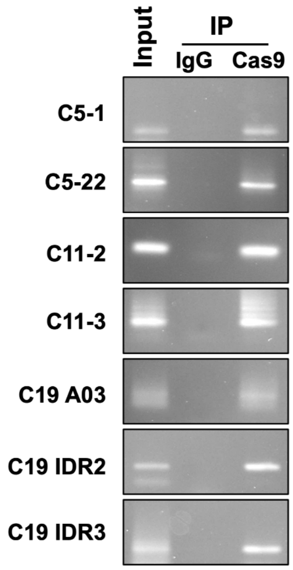


**Supplementary Figure 3**

**Detection of dCas9/sgRNA bound DNA sequences by ChIP-PCR by ChIP-PCR in CRISPR-Sirius cells.** U2OS chromatin was fixed and fragmented. Immunoprecipitation (IP) was performed using a Cas9 antibody (IP-Cas9) or an IgG antibody as a negative control, alongside a total chromatin baseline sample (Input). PCR was then performed to evaluate the enrichment of dCas9/sgRNA at specific targeted chromosomal regions.

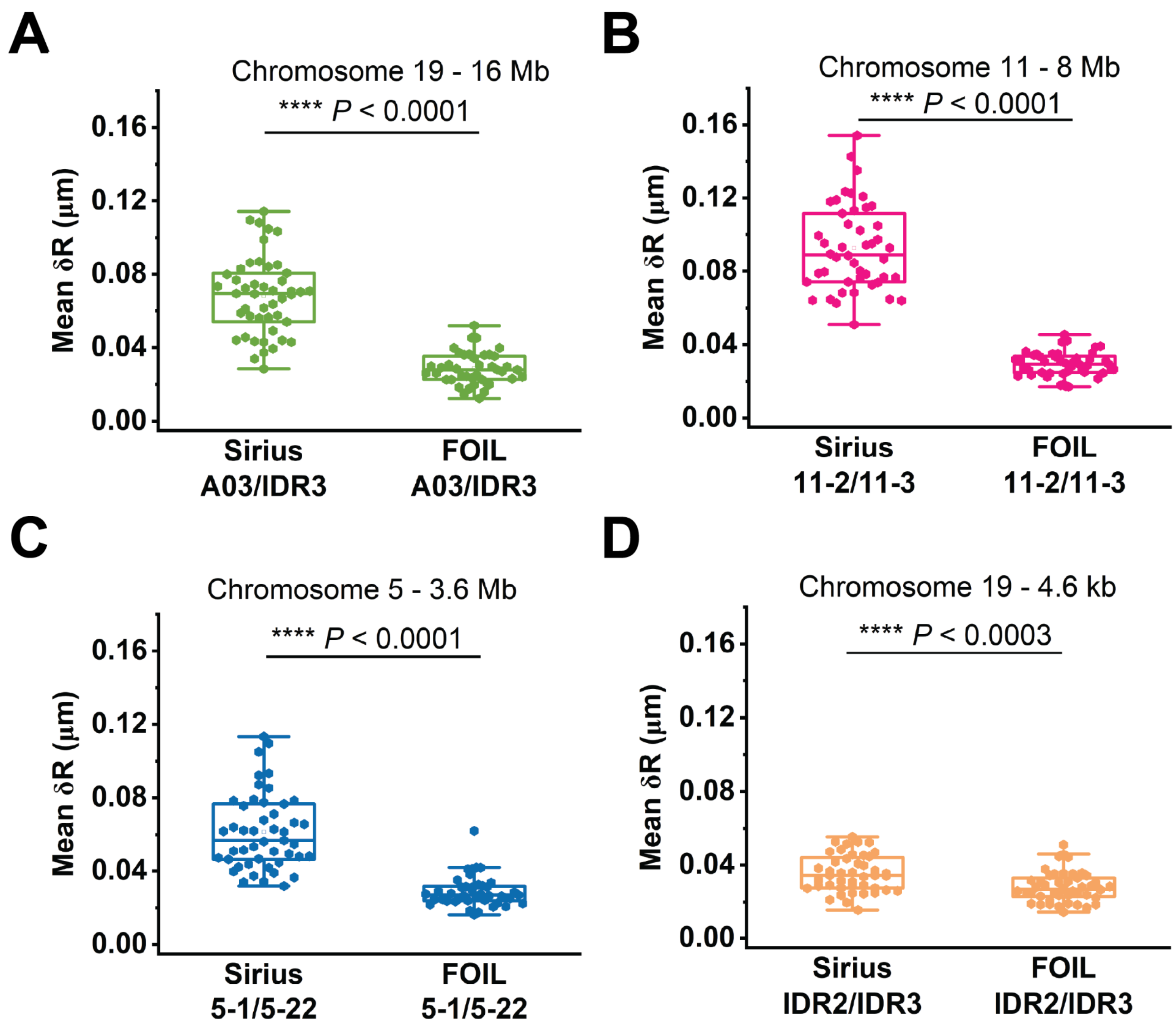


**Supplementary Figure 4**

**Temporal fluctuation of the distance (δR) between locus pairs.** Box plots show temporal fluctuation of the distance between locus pairs **(A)** C19-A03/C19-IDR3, **(B)** C11-2/C11-3, **(C)** C5-1/C5-22, and **(D)** C19-IDR2/C19-IDR3, labeled by either Sirius-sgRNA or FOIL-sgRNA. Imaging was performed over 30 frames at 500 ms intervals. $N_{C19\text{-}A03\text{-}C19\text{-}IDR3\text{-}Sirius}$=47, $N_{C19\text{-}A03\text{-}C19\text{-}IDR3\text{-}FOIL}$=41, $N_{C11\text{-}2C11\text{-}3\text{-}Sirius}$=46, $N_{C11\text{-}2C11\text{-}3\text{--}FOIL}$=45, $N_{C5\text{-}1C5\text{-}22\text{--}Sirius}$=47, $N_{C5\text{-}1C5\text{-}22\text{--}FOIL}$=41, $N_{C19\text{-}IDR2\text{-}C19\text{-}IDR3\text{-}FOIL}$=45, $N_{C19\text{-}IDR2\text{-}C19\text{-}IDR3\text{-}Sirius}$=44.

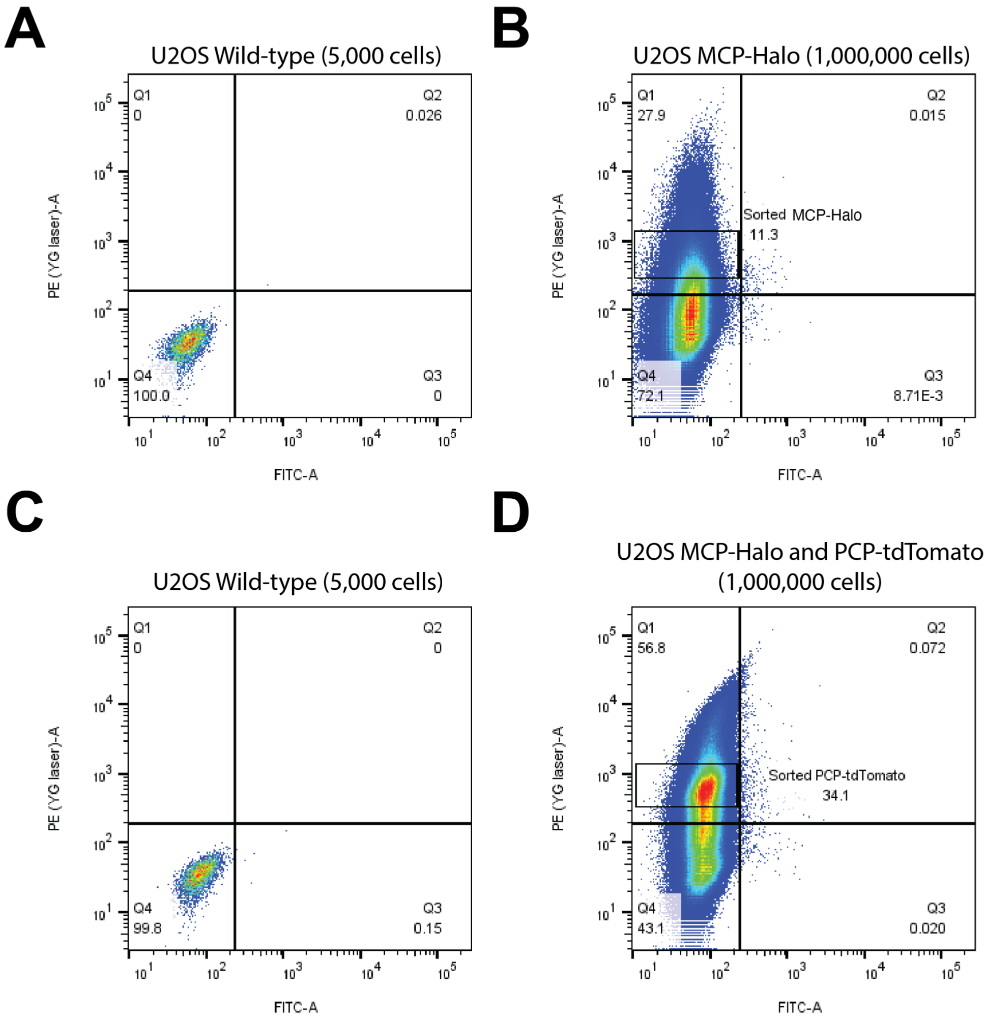


**Supplementary Figure 5**

**FACS gating strategy for generating the U2OS double red reporter cell line.** (A) - (B) Wild-type U2OS cells were transduced with MCP-Halo and sorted for HaloTag-JF549 signal. The gating box in (B) indicates the collected MCP-Halo positive population (~11.3%). (C ) -(D) U2OS$^{dCas9\text{-}HSA/MCP\text{-}HaloTag}$ cells were transduced with PCP-tdTomato and sorted for tfTomato fluorescence without HaloTag-JF549 (D) to isolate the double-labeled reporter line, , U2OS$^{dCas9\text{-}HSA/PCP\text{-}tdTOMATO/MCP\text{-}HaloTag}$. The gating box in (D) indicates the collected PCP-tdTomato positive population (~34.1%).

**Supplementary Table 1**
**CRISPR FOIL-gRNA sequences**

| | Chr. | Start | End | Targeting Sequences | Copies |
|---|---|---|---|---|---|
| 1 | Nontargeting (NT) | NA | NA | ACCG | NA |
| 2 | C5-1 | 83270 | 85599 | CGATGAGAGGCCnGG | 32 |
| 3 | C5-22 | 3637250 | 3639428 | TGTGTCTTGCACnGG | 50 |
| 4 | C11-2 | 60866662 | 60869195 | CACACATCAACAnGG | 68 |
| 5 | C11-3 | 68835708 | 68837303 | CTAGGACTCAGAnGG | 34 |
| 6 | C19-IDR2 | 376675 | 377633 | CCnCCCAGCCTCTCC | 36 |
| 7 | C19-IDR3 | 380777 | 382710 | AGCAGATGTAGGnGG | 45 |
| 8 | C19-A03 | 16042336 | 16044890 | AATGAGTGAGCGnGG | 29 |
| 9 | C19-C19q | 39200000 | 56100000 | TCCCTCAGACCCnGG | 836 |

**Supplementary Table 2**
**Vectors for CRISPR-FOIL**

| | Vector Name | RNA aptamers | Fluorescence | Addgene number |
|---|---|---|---|---|
| 1 | pPUR-hU6-sgRNA-Sirius-8XMS2 | 8XMS2 | - | 121942 |
| 2 | pPUR-mU6-sgRNA-Sirius-8XPP7 | 8XPP7 | - | 121943 |
| 3 | pHAGE-EFS-MCP-HALOnls | - | Red: JF549 | 121937 |
| 4 | pHAGE-EFS-PCP-GFPnls | - | Green: GFP | 121938 |
| 5 | pPUR-CMV-FOIL1-gRNA-8XMS2-2XBSL-FOIL2-gRNA-8XPP7 | 8XMS2, 8XPP7 | - | - |
| 6 | pHAGE-EFS-PCP-tdTOMATOnls | - | Red: tdTomato | - |

**pHAGE-EFS-PCP-NLSb-tdTOMATOnls sequence**

EFS: EF-1 Alpha Short promoter
PCP
NLSb: bipartite nuclear localization signal (from XRRC1)
tdTOMATO: codon optimized tandem tomato
nls: SV40 nuclear localization signal

tcgagtggctccggtgcccgtcagtgggcagagcgcacatcgcccacagtccccgagaagttggggggaggggtcggcaattgaaccggtgccta
gagaaggtggcgcggggtaaactgggaaagtgatgtcgtgtactggctccgcctttttcccgagggtgggggagaaccgtatataagtgcagtagtcg
ccgtgaacgttctttttcgcaacgggtttgccgccagaacacaggtgtcgtgacgcggttccggttaaggtgaacgcgtgcaggctggcgccaccatg
ggttccaaaaccatcgttctttcggtcggcgaggctactcgcactctgactgagatccagtccaccgcagaccgtcagatcttcgaagagaaggtcggg
cctctggtgggtcggctgcgcctcacggcttcgctccgtcaaaacggagccaagaccgcgtatcgcgtcaacctaaaactggatcaggcggacgtcg
ttgattccggacttccgaaagtgcgctacactcaggtatggtcgcacgacgtgacaatcgttgcgaatagcaccgaggcctcgcgcaaatcgttgtacg
atttgaccaagtccctcgtcgcgacctcgcaggtcgaagatcttgtcgtcaaccttgtgccgctgggccgtggtggcggagggactagtTCCCCT
AAAGGTAAACGGAAGTTGGATTTGAACCAAGAAGAAAAAAAGACCCCGAGTAAGCCACCG
GCACAGCTTTCCCCCTCCGTGCCTAAGCGGCCTAAACTTCCCgcggccgcgccgtggcctccggtagccatat
ggtgagcaagggcgaggaggtgattaaagaatttatgcgctttaaagtgagaatggaaggatccatgaatggacacgaatttgaaattgaaggagaag
gagaaggacgcccttatgaaggaacacagacagccaaactgaaagtgacaaaaggaggacctctgccttttgcctgggatattctgtcccctcagttta
tgtatggatccaaagcctatgtgaaacaccctgccgatattcctgattataaaaaactgtcctttcctgaaggatttaaatgggaaagagtcatgaattttga
agatggaggactggtgacagtgacacaggattcctccctgcaagatggaacactgatttataaagtgaaaatgagaggaacaaattttcctcctgatgg
acctgtcatgcagaaaaaaacaatgggatgggaagcctccacagaacgcctctatcctcgcgatggagtgctgaaaggagaaattcatcaggccctc
aaactgaaagatggaggacactatctggtcgaatttaaaacaatttatatggccaaaaaacctgtgcagctccctggatattattatgtggatacaaaactg
gatattacatcccacaatgaagattatacaattgtggaacagtatgaacgctccgaaggacgccatcatctgtttctggctagcgggcatggcaccggca
gcaccggcagcggcagctccggcaccgcctcctccgaggacaacaacatggccgtcatcaaagagttcatgcgcttcaaggtgcgcatggagggct
ccatgaacggccacgagttcgagatcgagggcgagggcgagggccgcccctacgagggcacccagaccgccaagctgaaggtgaccaagggc
ggcccccctgcccttcgcctgggacatcctgtcccccagttcatgtacggctccaaggcgtacgtgaagcaccccgccgacatccccgattacaagaa
gctgtccttccccgagggcttcaagtgggagcgcgtgatgaacttcgaggacggcggtctggtgaccgtgacccaggactcctccctgcaggacggc
acgctgatctacaaggtgaagatgcgcggcaccaacttcccccccgacggccccgtaatgcagaagaagaccatgggctgggaggcctccaccga
gcgcctgtaccccgcgacggcgtgctgaagggcgagatccaccaggccctgaagctgaaggacggcggccactacctggtggagttcaagacca
tctacatggccaagaagcccgtgcaactgcccggctactactacgtggacaccaagctggacatcacctcccacaacgaggactacaccatcgtgga
acagtacgagcgctccgagggccgccaccacctgttcctgtacggcatggacgagctgtaccgactcgagccaaagaaaaagcggaaagtgtaa
